# Serum Lewis fucosylation reports on survivorship of patients with septic shock upon admission to intensive care unit

**DOI:** 10.1101/2025.06.23.661221

**Authors:** The Huong Chau, Sayantani Chatterjee, Liam Caulfield, Anastasia Chernykh, Matthew Traini, Heeyoun Hwang, Rebeca Kawahara, Emily J. Meyer, David J. Torpy, Morten Thaysen-Andersen

## Abstract

Septic shock, the excessive immune response to pathogen infection, poses a major health concern accounting globally for ∼20% of all deaths. Current methods to establish disease severity are unacceptably slow, unspecific and insensitive, hindering timely and effective treatment. Aiming to identify easy-to-assay glyco-signatures that may identify and guide the clinical management of the most critically unwell patients, we applied quantitative glycomics and glycoproteomics to sera longitudinally collected from septic shock survivors (n=29) and non-survivors (n=8). Glycomics of all 134 serum samples (sampled daily until recovery/death) revealed significant *N*-glycome dynamics across both patient groups. Unsupervised clustering of the serum *N*-glycome upon ICU admission (day 1) indicated survivorship-specific glyco- signatures. We therefore employed machine learning to train a random forest model using the serum *N*-glycome data. The model accurately classified survivorship outcomes of 35 of 37 patients (specificity 94.6%) and correctly predicted 6 of 8 non-survivors (sensitivity 75%) based on ICU day 1 data. Further interrogation of the serum *N*-glycome data revealed elevated Lewis fucosylation in non-survivors relative to survivors at ICU admission, a finding recapitulated by glycoproteomics. Amongst 58 other serum proteins strongly linked to acute phase response and stress pathways, alpha-1-acid-glycoprotein was identified as a principal carrier of Lewis fucosylation with a potential to stratify septic shock survivors from non-survivors (AUC 0.935). This study lays a foundation for risk stratification of septic shock patients by uncovering easy-to-assay glyco-signatures that identify individuals with poor survival outcomes upon ICU admission, with the potential to translate to early individualised clinical care at the bedside.

## Introduction

Sepsis is the exaggerated immune response to pathogen infection and often escalates into life-threatening septic shock. These serious conditions pose a major health concern accounting globally for around 20% of all reported deaths costing annually a staggering $30 billion (USD) worldwide (1–5).

Despite the unacceptable burden of sepsis and septic shock on the health care systems, advances in early diagnosis, risk stratification, and tailored therapeutic interventions have been modest over the past two decades (6). A key challenge in the clinical management of septic shock is the lack of reliable biomarkers that can rapidly and accurately provide insight into the disease trajectory of individual patients. Current diagnostic approaches, including culture-based identification of pathogens and non-specific markers of inflammation, are often too slow or insensitive to support the timely decision-making required for such acute health conditions (7). Moreover, existing prognostic tools lack the specificity needed to guide personalised treatment pathways, particularly within the critical first 24 h of admission to the intensive care unit (ICU) when intervention is likely to benefit patients the most (8). Despite the severity of sepsis and septic shock, there have been limited advances in patient management leaving clinicians with few, often imprecise, therapeutic and prognostic tools (9). Sepsis is a complex syndrome and remains poorly understood, governed by a dynamic interplay of competing pro- and anti-inflammatory processes that ultimately shape patient outcomes (10).

To address the critical lack of diagnostic and prognostic tools, novel biomarker strategies that can capture dynamic aspects of the host response to infection and reflect the health status are urgently needed. In this context, glycosylation, the covalent attachment of complex sugar moieties (glycans) to proteins, has emerged as a promising yet still underutilised source of biomarker information (11, 12). Glycosylation is a prevalent modification of proteins that is widely recognised to undergo considerable structural changes with altered physiology and across a wide spectrum of disease conditions (13–15) including those accompanying sepsis and septic shock events (16–21). Glycoproteins present in blood serum, such as alpha-1-acid glycoprotein (AGP1) and immunoglobulins, are key mediators of the acute phase response and are subject to tightly regulated glycan remodelling in response to systemic insults (22–25).

Serum, a readily accessible clinical biospecimen, is rich in glycoproteins and reflects the systemic physiology of patients (26). Importantly, serum is routinely collected from critically ill individuals in ICU settings from central venous catheters, making glycan-based diagnostics compatible with existing clinical routines. Advances in high-resolution mass spectrometry and data analytics now enable comprehensive profiling of serum glycosylation with high sensitivity, precision, and throughput (27–31). Notably, recent improvements in glycomics and glycoproteomics techniques have opened opportunities to quantitatively map glycans in complex mixtures providing information of their fine structural details as well as their protein carriers and specific polypeptide attachment sites, knowledge that can be critical to understand the molecular mechanisms of and establish markers for complex diseases such as septic shock (32–37).

Previous work by our group has shown that serum *N*-glycosylation is significantly altered in modestly ill patients with bacteraemia, defined as the presence of pathogenic bacteria in the blood stream (19). In that study, we found that the serum *N*-glycome holds a promise to stratify these pre-septic shock individuals based on the infecting bacterial pathogen. Expanding on these findings, we recently investigated severely ill septic shock patients infected by different infectious agents, and found prominent pathogen class-specific serum *N*-glycome alterations in particular for the subset of individuals experiencing candidemia, a serious bloodstream infection caused by the fungus Candida (38). These studies highlight the potential of glycan-based profiling to identify the disease-causing pathogen(s) and reveal pathogen-specific host responses that are mirrored in the serum *N*-glycome. However, the prognostic utility of serum glycosylation, particularly its capacity to early in the disease predict survivorship of critically ill patients with septic shock, remains largely unexplored.

Based on this rationale, we hypothesised that serum glycosylation signatures may provide prognostic insight into disease severity and survivorship outcome of patients with septic shock. This work contributes to the growing body of evidence supporting the clinical utility of serum glycoprofiling and addresses a significant unmet need in the precision management of individuals experiencing septic shock.

## Methods

### Serum collection

Sera from septic shock patients were obtained from whole blood (1-10 mL/drawing) collected with informed consent at the Royal Adelaide Hospital (RAH), Adelaide, Australia. Specifically, sera were collected from 37 patients clinically diagnosed with septic shock (mean baseline SOFA score of 11 (range 4-19) and APACHE score of 23 (range 9-41)) immediately upon admission to the ICU at RAH (day 1) and then daily until recovery (ICU discharge) or death, see **Supplementary Table S1** for details of the patient cohort and key patient metadata. From this cohort, 8 patients died within ICU care from whom a total of 40 serum samples were longitudinally collected (average 5 samples/patient), while 29 patients recovered and were discharged from the ICU from whom 94 serum samples were longitudinally collected (average 4 samples/patient). Sera were aliquoted and stored at -20°C until use. Ethics approval for the collection and biochemical analysis of the septic shock sera was obtained from the RAH Human Research Ethics Committee (HREC/16/RAH/29).

### Protein handling

The protein concentrations of the neat serum samples were determined using a bicinchoninic acid assay (Thermo Fisher Scientific), following the manufacturer’s protocol and using a bovine serum albumin standard curve. Samples were diluted with triethylammonium bicarbonate (TEAB) to a final concentration of 1.5 µg/µL serum protein in 50 mM TEAB. Samples were reduced with 10 mM dithiothreitol (DTT) for 45 min at 56°C, alkylated with 30 mM aqueous iodoacetamide (both final concentration) for 30 min in the dark and quenched with excess DTT. The prepared serum protein extracts were used for both glycomics and glyco/proteomics as described below.

### *N*-glycomics sample preparation

Glycomics sample preparation of serum protein extracts followed a well-established method (39–41). In brief, 50 µg of reduced and alkylated proteins per sample was spotted onto an activated 0.45 μm PVDF membrane (Merck-Millipore), dried, stained with Direct Blue, and excised. The excised spots were transferred to separate wells in a flat bottom polypropylene 96-well plate (Corning Life Sciences, Melbourne, Australia), blocked with 1% (w/v) polyvinylpyrrolidone in 50% (v/v) aqueous methanol, and washed with ultra-pure water from a MilliQ source (used throughout the protocol).

The *N*-glycans were exhaustively released using 0.5 U/μL *Elizabethkingia miricola* peptide:*N*- glycosidase F (PNGase F) recombinantly expressed in *Escherichia coli* (Promega) per well and incubated for 16 h at 37°C. The released *N*-glycans were transferred into fresh tubes and hydroxylated by the addition of 100 mM aqueous ammonium acetate, pH 5 for 1 h at 20°C. The glycans were reduced using 1 M sodium borohydride in 50 mM aqueous potassium hydroxide for 3 h at 50°C. The reduction reaction was quenched using glacial acetic acid.

Dual desalting of the reduced *N*-glycans was performed using firstly strong cation exchange resin (AG 50W-X8 Resin, Bio-Rad), followed by porous graphitised carbon (PGC) resin custom packed as micro-columns on top of C18 discs (Merck-Millipore) in P10 solid-phase extraction (SPE) formats. Following micro-column equilibration and sample loading and washing, the *N*-glycans were eluted from the PGC-C18-SPE micro-columns using 0.1% trifluoroacetic acid (TFA)/50% acetonitrile (ACN)/49.9% water (v/v/v), dried and resuspended in 20 µL water. Samples were spun at 14,000 × *g* for 10 min at 4°C and the clear supernatant fractions were carefully transferred to high recovery glass vials (Thermo Fisher Scientific) to avoid debris and particulates in the LC-MS/MS injection vials.

### *N*-glycan profiling

The *N*-glycans were profiled using a well-established PGC-LC-MS/MS method (19, 38, 42, 43). In brief, the *N*-glycan samples were injected on a HyperCarb KAPPA PGC-LC column (particle/pore size, 3 µm/250 Å; column length, 30 mm; inner diameter, 1 mm, Thermo Hypersil, Runcorn, UK) heated to 50°C. To avoid data acquisition batch effects and injection sequence bias, the 134 *N*-glycan samples were injected in a randomised order considering both survivorship status and ICU time points. The *N*-glycans were separated over the following 60 min gradient of solvent B (70% ACN in 10 mM ammonium bicarbonate) in solvent A (10 mM aqueous ammonium bicarbonate). The gradient parameters were: 0 to 3 min - 0% B, 4 min - 14% B, 40 min - 40% B, 48 min - 56% B, 50 to 54 min - 100% B, 56 to 60 min - 0% B. The gradient was delivered by a 1260 Infinity Capillary HPLC system (Agilent) operating with a constant flow rate of 20 µL/min. The separated *N*-glycans were introduced directly into the mass spectrometer, ionised using electrospray ionisation and detected in negative ion polarity mode using a linear trap quadrupole Velos Pro ion trap mass spectrometer (Thermo Fisher Scientific). The acquisition settings included a full MS1 scan acquisition range of *m/z* 570– 2,000, resolution of *m/z* 0.25 full width half maximum and a source voltage of +3.2 kV. The automatic gain control (AGC) for the MS1 scans was set to 5 × 10^4^ with a maximum accumulation time of 50 ms. For the MS/MS events, the resolution was set to *m/z* 0.25 full width half maximum, the AGC was 2 × 10^4^ and the maximum accumulation time was 300 ms. Data-dependent acquisition was enabled for all samples. The three most abundant precursors in each MS1 full scan were selected for fragmentation using resonance activation (ion trap) collision-induced dissociation (CID) at a normalised collision energy (NCE) of 33%. Dynamic exclusion of precursors was inactivated. All MS and MS/MS data were acquired in profile mode. The mass accuracy of the precursor and product ions was typically better than 0.2 Da. The LC-MS/MS instrument was tuned and calibrated, and its performance bench marked using well-characterised bovine fetuin *N*-glycan standards analysed at the same time as the samples of interest.

### *N*-glycomics data analysis

The generated LC-MS/MS raw data files (made publicly available via GlycoPOST (44), https://glycopost.glycosmos.org/preview/177519572068520a17e26ef, accession number GPST000599, password: 4352) were browsed, interrogated, and manually annotated using Xcalibur v2.2 (Thermo Fisher Scientific), GlycoMod (45) and GlycoWorkBench v2.1 (46) as previously described (19). Briefly, glycans were identified based on the match between the observed and theoretical monoisotopic precursor mass and MS/MS fragmentation pattern generated *in silico* using GlycoWorkBench, and the expected relative and absolute PGC-LC retention time of each glycan (see **Supplementary Table S2** for all tabulated *N*-glycome data). The relative abundances of the confidently identified *N*-glycans were determined from area-under-the-curve (AUC) measurements based on extracted ion chromatograms performed for all relevant charge states of the monoisotopic precursor *m/z* using RawMeat v2.1 (Vast Scientific) and Skyline (64-bit) v24.1 (47, 48).

A machine learning model was established to assess whether the serum *N*-glycome profiles measured upon ICU admission (day 1) predict survivorship outcomes of the septic shock patients. For this, a supervised random forest (RF) model was established using *N*-glycomics data acquired from the longitudinally collected serum samples. To ensure independence of the training and test data and maximise the clinical relevance, the model was trained exclusively on serum *N*-glycome data collected from ICU day ≥ 2 and tested using ICU day 1 data. All machine learning modelling, including data pre-processing, feature selection, model training, and evaluation, was performed in R using custom scripts. Quantitative non-transformed glycome data were used as input into the ML model. Sensitivity and specificity were used to assess the performance of the model to predict patient survivorship.

### *N*-glycoproteomics and proteomics sample preparation

For glyco/proteomics, the serum protein extracts from ICU day 1 (37 samples) were exhaustively digested using sequencing-grade modified porcine trypsin (Promega) at a 1:50 ratio (enzyme:substrate, w/w) at 37°C for 17 h. The digestion was stopped by acidification with 1% (v/v) TFA (final concentration). Serum peptides were desalted using Oligo R3-C18-SPE clean up as previously described (32, 33). The desalted peptides were split into i) a minor fraction (10%) used for proteome analysis by direct LC-MS/MS without glycopeptide enrichment and ii) a major fraction (90%) used for glycoproteome analysis by LC-MS/MS after glycopeptide enrichment. For i), the fraction was dried and resuspended in 20 µL 0.1% (v/v) formic acid in preparation for direct LC-MS/MS. For ii), the fraction was dried and resuspended in 50 µL 0.1% (v/v) TFA in 80% (v/v) ACN in preparation for glycopeptide enrichment.

Glycopeptides were enriched utilising an established hydrophilic interaction liquid chromatography (HILIC)-SPE based enrichment method (49). Custom-made HILIC-C8-SPE micro-columns were prepared by inserting 1 mm Empore discs (Octyl C8 47 mm Extraction DISK 66882-U, Thermo Fisher Scientific) into P10 pipette tips (Eppendorf) onto which micro- columns of zwitterionic HILIC resin (ZIC-HILIC, 10 μm particle size, 200 Å pore size, kindly donated by Merck KGaA, Darmstadt, Germany) were packed (32, 33). The HILIC-C8-SPE micro-columns were conditioned after which the serum peptides were gently loaded and reloaded onto the micro-columns. Following thorough washing, glycopeptides were eluted sequentially using 50 µL 0.1% (v/v) TFA, 50 µL 25 mM aqueous NH4HCO3 and finally 50 µL 50% (v/v) ACN. The eluted glycopeptide fractions were combined, dried and resuspended in 20 µL 0.1% (v/v) formic acid for LC-MS/MS analysis.

### LC-MS/MS-based glyco/proteomics

Unenriched serum peptides were analysed in a single batch on a Q-Exactive HF-X Hybrid Quadrupole-Orbitrap mass spectrometer (Thermo Fisher Scientific) operated in positive ion polarity mode. To avoid data acquisition batch effects and injection sequence bias, samples were injected in a randomised order. Peptides (900 ng/sample) were loaded on to an Acclaim™ PepMap™ C18 reversed phase HPLC trap column (5 mm length × 300 μm inner diameter) and were separated on a nanoLC column (30 cm length x 75 μm inner diameter) packed in-house with Solidcore Halo® 2.7 μm 150 Å ES-C18 resin (Advanced Material Technology) operated at 45°C by an UltiMate™ NCS-3500RS HPLC system (Thermo Fisher Scientific). The HPLC produced a constant flow rate of 300 nL/min using a binary mobile phase system comprising 0.1% (v/v) formic acid (solvent A) and 0.1% (v/v) formic acid in 99% (v/v) ACN (solvent B). Following an initial equilibration for 5 min using solvent A, peptides were separated over a linear gradient ramping from 3.2-36% solvent B over 56 min followed by 36-95% solvent B over 2 min and an 8 min wash at 95% solvent B for a total run time of 75 min per sample. MS1 full scans (*m/z* 350-1,450) were acquired with a resolution of 60,000 and a normalised AGC target of 3 x 10^6^, a maximum injection time of 50 ms and a 1 s cycle time. Data-dependent acquisition was used to select the ten most abundant precursors in each MS1 scan for MS/MS, which were performed using higher-energy collisional dissociation (HCD) at 27.5% NCE. The precursor isolation window was *m/z* 1.4 and the MS/MS AGC target was 1 x 10^5^ with a resolution of 15,000. Selected precursors were dynamically excluded for 15 s. Unassigned and singly and highly charged precursors (z > 7) were excluded for MS/MS.

Enriched serum glycopeptides were analysed in a single batch on a Orbitrap Exploris 240 mass spectrometer (Thermo Fisher Scientific) operated in positive ion polarity mode. These samples were also injected in a randomised order. Glycopeptides (900 ng/sample) were loaded on to an Aurora Ultimate nanoLC column (25 cm length × 75 μm inner diameter, 1.7 μm particle size) operated at 60°C by a Vanquish Neo UHPLC System (Thermo Fisher Scientific). The HPLC provided a constant flow rate of 300 nL/min using the same solvent A and B as above. Peptides were separated over a linear gradient ramping from 3-35% solvent B over 90 min, 35-50% solvent B over 8 min followed by 50-90% solvent B over 2 min and a 10 min wash at 95% solvent B for a total run time of 110 min per sample. MS1 full scans (*m/z* 700-2,000) were acquired with a resolution of 120,000 using a standard AGC target with 100 ms maximum accumulation time. Data-dependent acquisition with a 3 s cycle time was used to select the ten most abundant precursors in each MS1 scan for MS/MS using HCD at 35% NCE. Fragment spectra were acquired within the orbitrap with an isolation window of *m/z* 1.6 and resolution of 15,000. The maximum accumulation time was 250 ms with a dynamic exclusion window of 20 s after a single isolation/fragmentation of any given precursor. Unassigned, singly charged and highly charged precursors (z > 7) were excluded for MS/MS.

### Glyco/proteomics data analysis

The raw glyco/proteomics LC-MS/MS datasets (deposited to the ProteomeXchange Consortium via the PRIDE partner repository (50), accession number PXD065024 (Username, Password: uLjWxdgjwRSE) were browsed and manually inspected using Xcalibur v2.2 (Thermo Fisher Scientific). Proteomics and glycoproteomics data from the unenriched and enriched peptide samples were analysed separately to identify and quantify the non-modified and glycosylated peptides and their source proteins, respectively. As an initial high-level analysis, the proportion of fragment spectra featuring the diagnostic marker for Lewis fucosylation (*m/z* 512.19 ± 2 ppm) was quantified across all LC-MS/MS files using an in-house script.

The glycoproteomics data (HCD-MS/MS) were searched using Byonic v4.5.2 (Protein Metrics) or the GPA search engine (51). For Byonic analyses, searches were performed employing a protein database comprising a reviewed UniProtKB *Homo sapiens* database (downloaded November 2021, 20,360 entries) alongside a predefined glycan search space of 310 mammalian *N-*glycans without sodium adducts. The search parameters were: precursor and product tolerance: 10 ppm and 20 ppm, respectively; semi-specific tryptic (RK) search alongside with N-ragged cleavage sites allowing 2 missed cleavages. Methionine oxidation (+15.994915 Da @ M) was considered a common variable modification while cysteine carbamidomethylation (+57.021464 Da @ C) was considered a fixed modification. A maximum of two common and one rare (glycan) modification was permitted for each peptide candidate. A second set of searches were performed which targeted only those fragment spectra that contained a diagnostic ion for Lewis fucosylation at *m/z* 512.190 (representing Hex-(Fuc)- HexNAc [H+]) with an *m/z* tolerance of 0.02 Da and a rank cutoff of 50. With the addition of this search parameter only identifications that possessed this diagnostic ion would be included in the results. The output from all glycopeptide searches was filtered to < 1% false discovery rate at the protein level and 0% at the peptide level using a decoy database and only confidently identified glycopeptides (PEP 2D < 0.001) were considered. For GPA searches, the LC-MS/MS raw files were converted into mzML files using RawConverter (version 1.2.0.1; The Scripps Research Institute, USA) with peak picking (vendor MS1) and default options. MS1 and MS/MS files were then extracted from the mzML files with an in-house program coded by Python v3.8 and GPA (v2.0) searches were performed to classify the fucosylation type (core or Lewis) using an artificial neural network model generated with Tensorflow (v2.0) (51). The detailed parameters for the identification of N-glycopeptides were as following: noise peak thresholds of 50.0 and 2.0 for MS and MS/MS, respectively; precursor mass tolerance of ± 0.05 Da; MS/MS tolerances of ± 0.02 Da for HCD; The cut-off threshold of M−, and S− score as 1.2, and 98.0, respectively; less than 1.0 % of the estimated FDR; retention time window of 5.0 min. For all search strategies using both Byonic and GPA, the relative abundances of the identified *N-*glycopeptides or select subsets thereof were determined using a spectral counting approach considering the glycopeptide-to-spectrum matches (glycoPSMs).

For protein identification and quantification, the unenriched raw files were imported into MaxQuant v2.4.13.0. The Andromeda search engine was used to search the HCD-MS/MS data against the reviewed UniProtKB *Homo sapiens* database (downloaded November 2021, 20,360 entries) with a precursor and product ion mass tolerance of 4.5 ppm and 20 ppm, respectively. Carbamidomethylation of cysteine (57.021 Da) was set as a fixed modification. Oxidation of methionine (15.994 Da) and protein N-terminal acetylation (42.010 Da) were selected as variable modifications. All identifications were filtered to < 1% false discovery rate (FDR) at the protein and peptide level using a conventional decoy approach. For label-free AUC-based quantification, the ‘match between runs’ feature of MaxQuant was enabled with a 0.7 min match time window and 20 min alignment time window. Protein abundance was calculated based on the normalised protein intensity (LFQ intensity).

Gene ontology (GO) analysis of biological process was performed using the GO Enrichment Analysis tool (52, 53), with the protein carriers of Lewis fucosylation as input and using the human proteome as the reference set. GO terms with *p* < 0.01 were considered significantly enriched.

### Experimental design and statistical rationale

The discovery phase of the study was performed using serum longitudinally collected from a cohort of septic shock survivors (n = 29) and non-survivors (n = 8), which provides sufficient statistical power to detect glycosylation changes and associations to survivorship outcome. The quantitative glycomics and glycoproteomics data were subjected to different statistical tests performed without data transformation and were visualised in various ways with or without transformation of the relative abundance data. Unpaired or paired two-tailed Student’s t-tests were generally performed to compare glycosylation signatures either between or within the two patient groups where *p* < 0.05 was chosen as the confidence threshold. Heat-maps and hierarchical clustering analyses were performed with Perseus v1.6.7 using Euclidean distance with average linkages (54). The relative abundance values of the glycans were used as input data for these analyses after log2 transformation. For the targeted (hypothesis-driven) statistical analyses of glycans unpaired or paired one-tailed Student’s t-tests were performed to compare glycosylation signatures between or within the two patient groups where *p* < 0.05 was chosen as the confidence threshold. Receiver operating characteristic (ROC) curves of both univariate and multivariate analyses were performed using linear support vector machines classification methods. The confidence threshold was AUC > 0.75 with 1.00 representing perfect separation between patient groups. For volcano plot and the ROC curves that were performed using MetaboAnalyst (55), untransformed or log2 transformed relative abundance values of the glycans from each patient group were used as input data. If not mentioned otherwise, data have been plotted as mean (average) and error bars indicate SD. Replicates (biological/technical as indicated) and statistical significance have been stated in the figure legends.

## Results

### Leveraging integrated -omics methods to map the serum glycoproteome in septic shock

Driven by our goal to identify clinically informative glyco-signatures that may identify and thus guide the management of the most severely affected individuals with septic shock, we took a multi-omics approach to comprehensively survey the *N*-glycoproteome of serum collected from a valuable cohort of patients clinically diagnosed with septic shock that either recovered (n = 29 survivors) or died (n = 8 non-survivors) from the condition, **Figure 1A**. Importantly, serum was collected daily from both the survivors (from ICU admission to discharge, 2-6 samples per patient, in total n = 94 samples) and from the non-survivors (from ICU admission to death, 3-7 samples per patient, in total n = 40 samples). Different bacteria were identified as the disease-causing pathogens for most of the septic shock patients including *Pseudomonas aeruginosa* (n = 6), *Escherichia coli* (n = 12), *Staphylococcus aureus* (n = 10), *Prevotella melaninogenica* (n = 1) while no infecting pathogen was confidently identified for a subset of the septic shock patients (n = 8). The patient cohort included both males and females and spanned a large range in terms of age (40-86 years old) and body mass index (17.9-62.3 kg/m^2^), **Supplementary Table S1**. While the two patient groups were appropriately gender and age matched, the patients therefore formed a relatively heterogenous cohort reflecting the makeup of the individuals affected by this disease.

**Figure 1.**
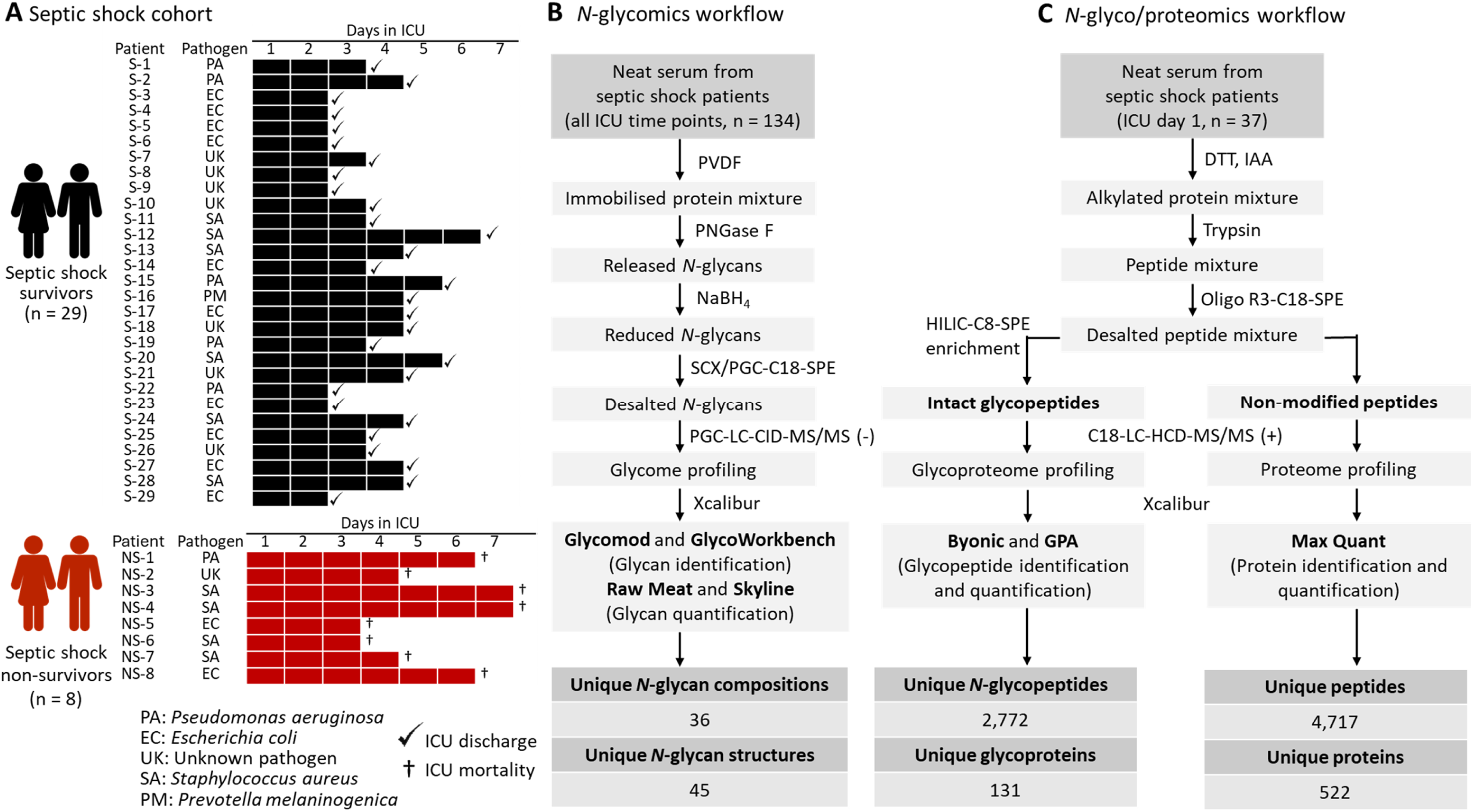
Multi-omics profiling of the serum N-glycoproteome during septic shock. **A**. Overview of the septic shock cohort comprising both patients who recovered (survivors, n = 29, labelled S-1 to S-29, in black) and patients who died (non-survivors, n = 8, NS-1 to NS-8, in red) from the disease. Serum was collected daily from ICU admission (day 1) until ICU discharge (√) or death (†) as indicated for each patient. A variety of disease-causing pathogens were identified across the patient cohort, see key. See **Supplementary Table S1** for additional metadata (age, gender, body mass index, baseline APACHE and SOFA scores) for the septic shock cohort. Experimental approach for the quantitative glycomics (**B**) and glyco/proteomics (**C**) applied to septic shock sera. The number of unique glycans, glyco/peptides and glyco/proteins identified in the septic shock sera are indicated.

Enabled by our glycomics-guided glycoproteomics technology (32, 33), we performed quantitative glycomics, glycoproteomics and proteomics of the septic shock serum samples, **Figure 1B**. Glycomics, involving the detailed profiling of detached *N*-glycans from all ICU sampling points (134 serum samples) using PGC-LC-MS/MS, quantitative mapped a total 45 *N*-glycan structures (including isomers) across 36 different glycan compositions within the septic shock serum *N*-glycome (detailed further below). To obtain important complementary information, the serum glycoproteome of the clinically valuable ICU day 1 sampling point (37 serum samples) was surveyed using glycoproteomics and proteomics involving reversed phased-LC-MS/MS of serum peptide mixtures that were either enriched or unenriched for intact glycopeptides, respectively. The glycoproteomics experiments identified a total of 2,772 unique (non-redundant) *N*-glycopeptides originating from 131 serum glycoproteins while the proteomics experiments, carried out to establish the relative protein levels across the serum samples, identified 4,717 unique peptides derived from 522 serum proteins. Collectively, these parallel analyses form, to the best of our knowledge, the, to date, most comprehensive map of the serum *N*-glycoproteome across the septic shock disease course.

### Surveying *N*-glycome dynamics in septic shock reveals signatures with stratification potential

The 45 *N*-glycan structures identified by the detailed PGC-LC-MS/MS-based glycomics profiling of septic shock sera featured, as expected from established serum *N*-glycome literature (19, 38, 56), primarily complex-type *N*-glycans spanning predominantly bi- and tri-antennary *N*-glycans and less abundant mono-antennary and bisecting GlcNAcylated *N*- glycans, **Figure 2A**. Also aligning with expectations, oligomannosidic- and hybrid-type *N*- glycans formed relatively minor subsets of the septic shock serum *N*-glycome, see **Supplementary Table S2** for all tabulated *N*-glycome data.

**Figure 2.**
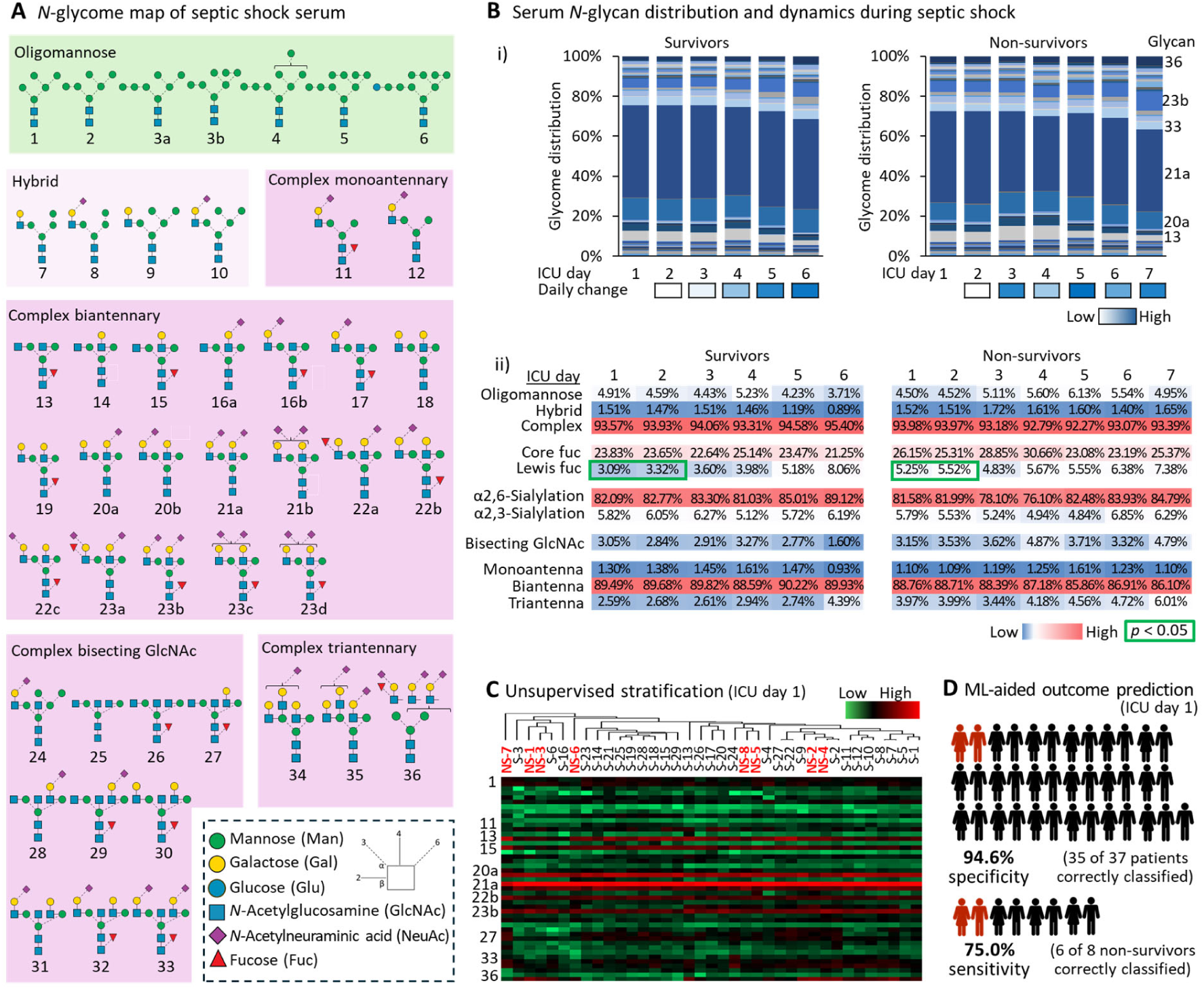
*N*-glycan profiling of septic shock sera reveals signatures with stratification potential. **A.** Map of *N*-glycan structures identified in septic shock sera. Glycan cartoons are drawn using the latest SNFG nomenclature (74), see insert for key. **B**. i) Distribution of serum *N*-glycans from survivors (left, average of n = 29) and non-survivors (right, average of n = 8) from the individual ICU sampling point during a septic shock event. Main glycan structures are labelled. See **Supplementary Table S2** for all tabulated *N*-glycome data. The relative day-to-day change in serum *N*-glycome (average across all patients) is indicated by heat map colours ranging from low (white) to high (blue) dynamics, see key. ii) Longitudinal plot of the relative level of discrete glycan features in sera from septic shock survivors (left, average of n = 29) and non-survivors (right, average of n = 8). Values are accompanied by heat map colours ranging from low (blue) to medium (white) to high (blue) relative levels. Two-tailed Student’s t-tests were used to compare glycan features across survivors and non-survivors for each ICU time point; green box indicates statistical significance (*p* < 0.05). **C**. Partial stratification of septic shock non-survivors (NS, bolded in red) from survivors (S, in black) using unsupervised hierarchical clustering on the entire serum *N*- glycome data from ICU day 1. The *N*-glycome data were visualised with heat map colours ranging from low (green) to medium (black) to high (red) relative abundance, see key. **D**. ML-guided prediction of survivorship outcome of septic shock patients based on the entire serum *N*-glycome. A random forest model was trained using all serum *N*-glycome data from all ICU time points except ICU day 1 and the performance (prediction accuracy i.e. specificity and sensitivity) was tested separately using the clinically strategic ICU day 1 data.

While protein *N*-glycosylation is considered relatively stable in serum with limited intra- and inter-personal variations even under pathophysiological conditions (23, 57, 58), longitudinal plots of the *N*-glycan distribution revealed considerable dynamics of the serum *N*-glycome over the course of a septic shock event, **Figure 2Bi**. For septic shock survivors, the day-to-day *N*-glycome fluctuations paralleled recovery and was most pronounced upon discharge from the ICU whereas for the non-survivors profound fluctuations in the *N*-glycome were found throughout the mid-to-late disease stages mirroring the progression of sepsis shock to ICU death). Assessing the dynamics at the *N*-glycan feature level, the sera however appeared relatively stable throughout the septic shock event for both survivors and non-survivors, **Figure 2Bii**. As such, no significant differences in *N*-glycan type distribution (e.g. oligomannose, 3.7%-6.1%), core fucosylation (21.2%-30.7%), α2,3- (4.8%-6.9%) and α2,6-sialylation (76.1%-89.1%) and glycan branching (e.g. biantennary, 86.1%-90.2%) were observed within or between the patient groups while in ICU care. As an important exception to these trends, the outer arm (antennary, hereafter referred to as Lewis) fucosylation was found to be significantly raised in non-survivor sera on ICU day 1 and day 2 (5.25%-5.52%) relative to levels in survivor sera (3.09%-3.32%, *p* < 0.02-0.03). This survivorship-specific feature is explored in greater detail below.

Taking firstly an unsupervised approach, we used hierarchical clustering to explore if the entire *N*-glycome in serum collected upon ICU admission (ICU day 1) could discriminate septic shock survivors from non-survivors, **Figure 2C**. While no clean separation was achieved between the two patient groups, the unsupervised clustering analysis partially grouped the septic shock non-survivors indicating some stratification potential of the serum *N*-glycome at the early disease stage. We therefore sought to establish glycome-based prediction model using a machine learning (ML) approach employing all ICU time points except for ICU day 1 data to train a random forest model. The serum *N*-glycome data measured at ICU day 1 were then used to test the prediction accuracy of model. Encouragingly, the ML-guided model showed very high specificity (94.6%) by correctly predicting the survivorship outcome of 35 of the 37 septic shock patients as well as high sensitivity (75.0%) by correctly classifying 6 of the 8 non-survivors, **Figure 2D**. Upon inspection of the ML-guided prediction model, glycan 22a and glycan 23a (both featuring Lewis fucosylation, see below) as well as glycan 35, were found to be the main contributors to the survivorship outcome prediction.

### Lewis fucosylated glycans stratify non-survivors from survivors early in sepsis shock

Quantitative analysis of the individual *N*-glycan structures identified in septic shock serum revealed that glycan 22a and glycan 23a were significantly elevated in non-survivors relative to survivors, **Figure 3A**. No other *N*-glycan isomers appeared significantly different between the two patient groups. Detailed CID-MS/MS-based characterisation was used to elucidate the fine structural details of the observed serum *N*-glycans including glycan 22a and glycan 23a, **Figure 3B**. A common feature of glycan 22a and glycan 23a was the Lewis fucosylation on the 3’ arm of the biantennary complex-type *N*-glycan structures appearing as a Lewis epitope on glycan 22a (i) and a sialyl Lewis epitope on glycan 23a (ii). In line with robust literature on septic shock serum (20, 59, 60), our detailed PGC-LC-MS/MS-based glycomics data indicated that the glycoepitopes were Lewis x (Le^x^) and sialyl Lewis x (sLe^x^) rather than Le^a^ and sLe^a^. In addition to the Lewis fucosylation, the biosynthetically related glycan 22a and glycan 23a both featured a sialyl LacNAc on the 6’ arm. Using the longitudinally collected serum samples to map the temporal expression of glycan 22a and glycan 23a across the initial stages of a septic shock event (ICU day 1-4), showed that these two glycans were significantly over-represented in non-survivor relative to survivor sera obtained on ICU day 1 (glycan 22a: *p* = 0.0004, glycan 23a: *p* = 0.0076) and ICU day 2 (glycan 22a: *p* = 0.0031, glycan 23a: *p* = 0.0346), **Figure 3C**. Resulting from a relative increase of these two glycan isomers within the survivor group, similar levels of glycan 22a and glycan 23a were generally detected at days 3 and 4 in those patients with ongoing septic shock requiring ICU care (*p* ≥ 0.05,). Excitingly, the biggest abundance difference between non-survivors and survivors was observed on the clinically important ICU day 1 for both glycan 22a (*p* = 0.0004) and glycan 23a (*p* = 0.0076) when intervention options are most beneficial, **Figure 3D**. Finally, targeted ROC analysis demonstrated a potential of both glycan 22a (AUC: 0.862) and glycan 23a (AUC: 0.815) to predict septic shock survivorship outcome, **Figure 3E**.

**Figure 3.**
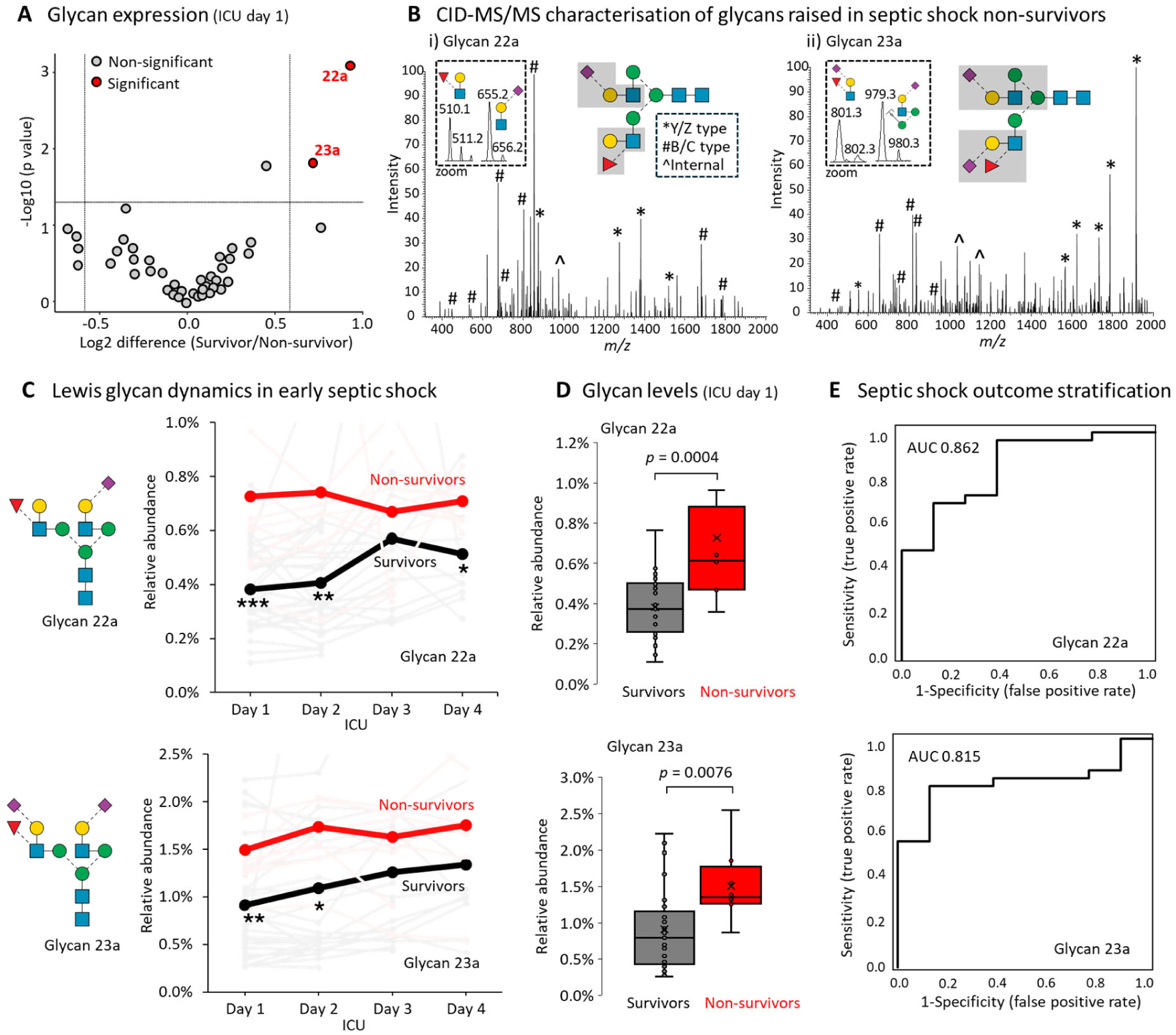
Lewis fucosylated glycans stratify non-survivors from survivors early in sepsis shock. **A.** Comparison of the relative abundance of individual serum *N*-glycans from survivors (n = 29) and non-survivors (n = 8). *N*-glycans that differ between patient groups (*p* < 0.05, glycan 22a and glycan 23a) are plotted in red; unpaired Student’s t-tests. **B**. CID-MS/MS-based characterisation of i) glycan 22a and ii) glycan 23a. Spectral features have been annotated (see insert for key to fragment ion types) with a focus on key terminal fragments including Lewis fucosylation-specific product ions. See **Supplementary Figure S1** for fully annotated MS/MS spectra. **C**. Longitudinal plots of glycan 22a (top) and glycan 23a (bottom) in sera from survivors (black trace, averages of 29 patients) and non-survivors (red trace, averages of 8 patients) in the early stages of septic shock (ICU day 1-4). Patient-specific traces are shown in faint grey (survivors) and faint red (non-survivors). Glycan levels were compared between patient groups at each ICU time point using unpaired Student’s t-tests, \**p* < 0.05, \*\**p* < 0.01, \*\*\**p* < 0.001. **D**. Boxplots showing relative level of glycan 22a (top) and glycan 23a (bottom) in sera collected on ICU day 1 from survivors (n = 29) and non-survivors (n = 8). Glycan levels were compared between patient groups using unpaired Student’s t-tests. **E**. ROC plots showing the ability of glycan 22a (top) and glycan 23a (bottom) measured on ICU day 1 to correctly predict the survivorship outcome of septic shock patients.

### Lewis fucosylated AGP-1 separates septic shock non-survivors from survivor on ICU day 1

Prompted by the glycomics data indicating a stratification potential of Lewis fucosylation in serum from early-stage septic shock, we performed comparative glycoproteomics of serum collected on ICU day 1 in attempts to quantitatively support this interesting observation and determine the protein carrier(s) of this outcome-specific glycosylation feature.

An initial high-level XIC-based monitoring of a Lewis fucosylation-specific fragment ion (*m/z* 512.19, representing Hex-(Fuc)-HexNAc [H+]) across the glycoproteomics LC-M/MS datasets indicated a visibly stronger signal trace in septic shock non-survivors relative to the trace strengths in survivors, **Figure 4Ai**. Quantifying the *m/z* 512 diagnostic ion at the MS/MS spectral level indeed showed a significantly higher level of Lewis fucosylation for non-survivors compared to survivors (*p* = 8.66 x 10^-5^), **Figure 4Aii** and **Supplementary Table S3**.

**Figure 4.**
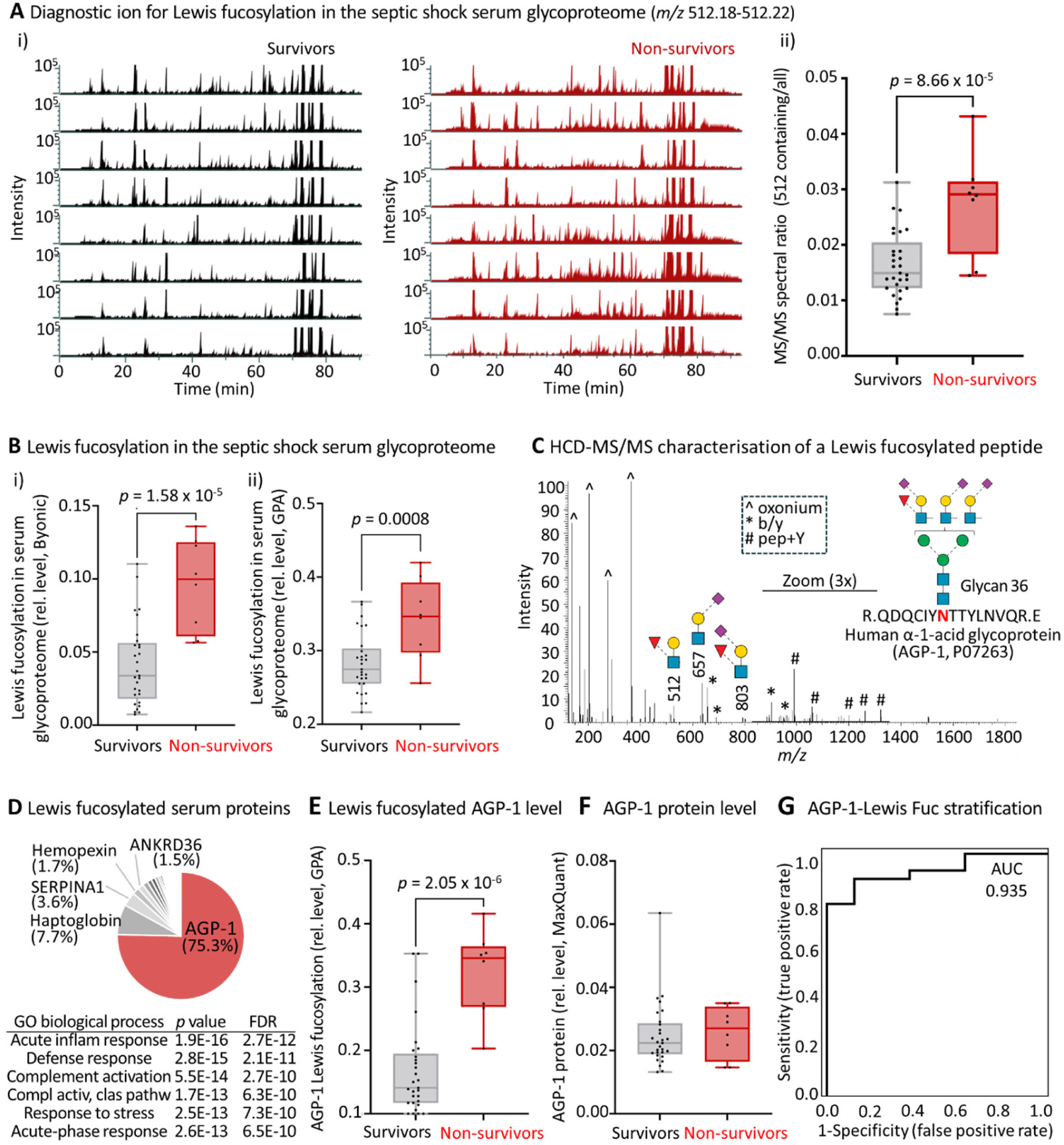
Lewis fucosylated AGP-1 separates septic shock non-survivors from survivors on ICU day 1. **A**. i) Stacked XICs tracking of a key diagnostic fragment ion specific for Lewis fucosylation (*m/z* 512.18-512.22) within LC-MS/MS-based glycoproteomics data of 8 septic shock survivors (black traces, left) and all 8 non-survivors (red traces, right). ii) Quantification of the relative level of Lewis fucosylation across patient groups as measured by the MS/MS spectral ratio (*m/z* 512 containing fragment spectra out of all MS/MS spectra, **Supplementary Table S3**) for all septic shock survivors (n = 29) and non-survivors (n = 8). Unpaired Student’s t-tests. **B**. Global level of Lewis fucosylation in the serum glycoproteome of septic shock survivors (n = 29) and non- survivors (n = 8) as determined using i) Byonic and ii) GPA search engines on LC-MS/MS-based glycoproteomics data from ICU day 1. See **Supplementary Table S4-S5** for all tabulated Byonic and GPA glycoproteome data, respectively, and **Supplementary Table S6** for relative Lewis fucosylation levels. Unpaired Student’s t-tests. **C**. HCD-MS/MS-based characterisation of a representative Lewis fucosylated glycopeptide derived from AGP-1. Spectral features have been annotated (see insert for key to fragment ion types) with a focus on key terminal fragments including Lewis fucosylation-specific product ions. An *m/z* region was magnified to visualise b/y peptide ions as indicated. **D**. Pie chart showing the Lewis fucosylated glycoproteins identified in septic shock serum. The five most abundant Lewis fucosylated proteins including AGP-1 (in red) have been labelled (see **Supplementary Table S7** for the entire list of identified Lewis fucosylated proteins in septic shock serum). Pathway enrichment analysis showing the six most enriched pathways related to the identified Lewis fucosylated glycoproteins in septic shock serum. **E**. Relative level of Lewis fucosylation carried by AGP-1 in sera from septic shock survivors (n = 29) and non-survivors (n = 8) as determined using GPA on LC-MS/MS-based glycoproteomics data. Unpaired Student’s t-test. See **Supplementary Table S8** for Lewis fucosylation data **F**. Relative level of AGP-1 protein in sera from septic shock survivors (n = 29) and non-survivors (n = 8) as determined using MaxQuant on LC-MS/MS-based proteomics (unenriched) data. See **Supplementary Table S9** for all tabulated proteome data and **Supplementary Table S10** for relative AGP-1 protein levels. Unpaired Student’s t-test. **G**. ROC plot showing the ability of Lewis fucosylated AGP-1 measured on ICU day 1 to accurately predict the survivorship outcome of septic shock patients.

Aiming to further support this observation, we analysed the glycoproteomics data using two different search engines, Byonic and GPA, with a focus on quantifying peptides carrying Lewis fucosylated, see **Supplementary Table S4** and **Supplementary Table S5** for all tabulated glycoproteome data from the Byonic and GPA search engines, respectively. On one hand, an innovative Byonic-based approach was used involving both a narrow (restricted) search against only those HCD-MS/MS spectra that contained a prominent *m/z* 512 ion diagnostic for Lewis fucosylation and an unrestricted search against all HCD-MS/MS spectra. Employing this approach, which allowed us to estimate the relative level of Lewis fucosylation in the serum glycoproteome, confirmed a raised level of Lewis fucosylation in septic shock non-survivors compared to survivors (*p* = 1.58 x 10^-5^), **Figure 4Bi** and **Supplementary Table S6**. On the other hand, we also employed the GPA search engine that has been trained to distinguish Lewis fucosylated glycopeptides from those carrying core fucosylation (51). Reassuringly, the GPA-based searches validated a higher level of Lewis fucosylation in the serum glycoproteome of non-survivors compared to survivors (*p* = 0.00080), **Figure 4Bii**. HCD-MS/MS-based characterisation of the peptides carrying Lewis fucosylated *N*-glycan structures not only revealed the protein carrier identities (in this representative example, AGP-1), but also confirmed the structure of the conjugated glycans (glycan 36) and the terminal glycoepitopes (sialyl Lewis fucosylation i.e. *m/z* 512 and 803 and sialyl LacNAc i.e. *m/z* 657), **Figure 4C**.

The Byonic-annotated glycoproteome data were then interrogated to identify the proteins carrying Lewis fucosylation in septic shock serum. The 309 unique (non-redundant) Lewis fucosylated glycopeptides that were identified in septic shock serum were found to map to a total of 59 different source proteins including most prominently AGP-1 (75.3%) while haptoglobin (7.7%), alpha-1-anti-trypsin (SERPINA1, 3.6%), hemopexin (1.7%), and ankyrin repeat domain-containing protein 36A (ANKRD36, 1.5%) amongst many other serum proteins were of lower abundance, **Figure 4D** and **Supplementary Table S7.** Interestingly, pathway enrichment analyses using gene ontology (GO) of the 59 Lewis fucosylated glycoproteins identified in septic shock sera against a human proteome reference set revealed strong over- representation of pathways known to be associated with septic shock including, for example, acute inflammatory response (*p* = 1.9 x 10^-16^, FDR 2.7 x 10^-12^), complement activation (*p* = 5.5 x 10^-14^, FDR 2.7 x 10^-10^), response to stress (*p* = 2.5 x 10^-13^, FDR 7.3 x 10^-10^) and acute-phase response (*p* = 2.6 x 10^-13^, FDR 6.5 x 10^-10^).

Given the profound dominance of AGP-1 amongst the other less abundant Lewis fucosylated protein carriers, we explored the relative level of Lewis fucosylated AGP-1 between the patient groups and found a significant elevation in septic shock non-survivors relative to the survivor group (*p* = 2.05 x 10^-6^), **Figure 4E** and **Supplementary Table S8**. As AGP-1 similar to many other Lewis fucosylated serum proteins are known to be a positive acute phase protein and thereby raised in circulation as a result of systemic infection, we sought to establish the relative AGP-1 protein level in the two patient groups. As measured using the quantitative proteomics data (see **Supplementary Table S9** for all tabulated proteome data), the AGP-1 level was found to be similar in non-survivors and survivors (p ≥ 0.05) suggesting a glycosylation- rather than a protein-based regulation underpinning the molecular differences between the patient groups, **Figure 4F** and **Supplementary Table S10**. Excitingly, ROC analysis demonstrated that Lewis fucosylated AGP-1 accurately predicts survivorship outcome in septic shock patients from serum collected upon ICU admission (day 1) (AUC: 0.935), **Figure 4G**.

## Discussion

Septic shock, the unbalanced and often exaggerated immune response to infection, remains a major cause of global mortality, yet tools for accurate patient risk stratification and prediction of disease trajectory are still unavailable to ICU clinicians. To address this significant clinical gap, we here applied an integrated glycomics and glycoproteomics method to 134 serum samples from septic shock survivors and non-survivors, collected daily within the ICU until recovery or death. These efforts enabled us to generate the, to date, most comprehensive map of the serum *N*-glycoproteome across the septic shock disease course.

Our -omics investigations revealed dynamic changes in the serum *N*-glycome and identified distinct glycosylation patterns between survivors and non-survivors as early as upon ICU admission (day 1). Using an ML-guided approach, a random forest model trained on the serum *N*-glycome profiles correctly predicted the survival outcome of 35 of 37 patients using ICU day 1 data. Lewis fucosylation, in the form of Le^x^ and sLe^x^ glycoepitopes, carried predominantly by serum AGP-1, was significantly elevated in non-survivors and was a strong predictor of poor outcome upon ICU admission (AUC 0.935). These findings suggest that specific glyco-signatures, easily measurable within neat serum, may serve as biomarkers to inform prognosis and guide personalised treatment strategies of patients early in septic shock.

In line with our findings, an early study used simple lectin and antibody-based electrophoretic methods to suggest that sLe^x^-containing AGP-1 glycoforms, rather than the AGP-1 protein level itself, is raised in septic shock non-survivors relative to survivors (59). Our multi-omics approach not only validates these early findings, but also adds important biochemical details (structural depth and robust quantitation) and offers additional insights into the dynamics of the serum *N*-glyco(proteo)me through the exploration of the longitudinally collected serum samples. Temporally, the most prominent glycoprofile differences between the two patient groups were observed upon ICU admission suggesting that these distinct molecular signatures can be harnessed for prediction of disease trajectory.

Lewis-type antigens have been previously associated with leukocyte adhesion, selectin binding, and immunosuppressive signalling in cancer (61–63) and inflammation (64), but their use as prognostic markers in acute critical illness remains underexplored. The functional implications of these changes in glycosylation of AGP in inflammation and how serum Lewis fucosylation may mechanistically link to poor septic shock outcome are as yet unknown. Providing possible clues to these associations, Le^x^-containing AGP-1 has been suggested to interfere with leukocyte extravasation by binding to adhesion molecules such as endothelial receptors in the selectin family (65). Additionally, *O*-glycans can also carry Lewis motifs (66), and it remains to be investigated whether AGP-1 or other glycoproteins contribute *O*-linked Lewis fucosylation in septic shock and if these also inhibit trans-epithelial leukocyte migration.

While we and others did not observe any protein-level differences in AGP-1 between septic shock survivors and non-survivors, this positive acute phase protein is known to be substantially raised in septic shock sufferers (∼1.6-1.7 mg/ml) relative to healthy donors (∼0.8 mg/mL) (59). The additional AGP-1 in septic shock sera is predictably from hepatic origins, however, since neutrophils are also known to produce, store and upon activation release AGP- 1 with different glycosylation features than liver AGP-1 (67), future efforts are required to explore if septic shock survivors and non-survivors carry AGP-1 from different tissue origins as this may explain the different glycosylation across these two patient groups. Biosynthetically, Lewis-type fucosylation is mediated by α1,3/4-fucosyltransferases such as FUT3, FUT5, and FUT6 (68), which may conceivably be differentially regulated in septic shock survivors and non-survivors due to differences in cytokine signalling, epigenetics, or hepatic (dys)function. If Lewis fucosylation is found to be mechanistically involved in septic shock outcome, these FUT enzymes may be potential targets as novel therapeutic intervention strategies against septic shock.

Reflecting both the considerable technical challenge of performing large multi-omics studies as well as the logistic challenge of recruiting critically ill septic shock individuals for daily serum sampling within the ICU, a limitation of this study was the modest cohort size and the substantial patient heterogeneity, including differences in age, gender, BMI, infection source, treatment regimens, and comorbidities. Given the emerging links between the serum *N*- glycome and age, gender and other physiological conditions (69, 70), these various factors may influence the glycosylation in septic shock serum independently of clinical trajectory, and could confound associations if not adequately controlled. Sepsis remains a highly complex and heterogeneous syndrome, and to fully establish the diagnostic and prognostic utility of glycan-based biomarkers, future studies will require larger, prospectively stratified cohorts. Given our recent findings that the infecting pathogen class impacts the host serum glycoprofile in both pre-septic shock (71) and septic shock conditions (38), the inclusion of patients with viral and fungal infections across varying severities, and longitudinal monitoring of patients (from the early stages of sepsis through progression to septic shock within the ICU, and extending through to either death or recovery into the post-ICU period), will serve to validate the survivorship outcome-specific glycan signatures observed herein and determine their temporal dynamics. Large patient cohorts will also enable more thorough training and thus stronger predictive power of the ML models employed in this study. Additionally, expanding the analytical scope to other glycan classes, such as *O*-glycans, may provide further insight into the host response and potentially reveal other outcome marker candidates.

From a translational perspective, the ability of Lewis fucosylation as a biologically relevant glycan motif to report on septic shock patient outcome is interesting particularly given the current lack of prognostic tools available to ICU clinicians. The fact Lewis fucosylation can be analytically measured by unsophisticated methods (antibodies, lectins) directly in crude matrices including serum highlights the potential of this glyco-signature as a dynamic and functional biomarker candidate for risk stratification of patients with septic shock. Offering even more predictive performance, AGP-1-specific Lewis fucosylation may represent a viable target for point-of-care risk stratification using combined antibody- and lectin-based readouts in array formats (72, 73) or even by high through-put LC-MS/MS (24, 70) from a few microliters of neat serum or blood from septic shock sufferers. Such approaches could enable rapid, minimally invasive patient outcome prediction in the ICU, supporting more personalised and timely management of septic shock ultimately improving patient survival.

## Data availability

Glycomics LC-MS/MS raw data were deposited to GlycoPOST (44), accession number GPST000599, URL: https://glycopost.glycosmos.org/preview/177519572068520a17e26ef, password: 4352). The glycoproteomics and proteomics LC-MS/MS raw data were deposited to the ProteomeXchange Consortium (http://proteomecentral.proteomexhange.org) via the PRIDE partner repository (50), accession number PXD065024 (Username, Password: uLjWxdgjwRSE).

## Supplemental data

This article contains supplemental data.

## Acknowledgments

We thank Joshua Fehring and Dr Liisa Kautto for providing assistance and support in this study.

## Funding and additional information

T.H.C. is supported by an International Research Training Program Scholarship from Macquarie University (20224231).

S.C. was supported by an International Research Training Program Scholarship from Macquarie University (2017152).

H.H. was supported by the Korea Basic Science Institute (C523400)

1. R. K. was supported by the Cancer Institute of New South Wales (ECF181259).

E.J.M. is the recipient of a Royal Adelaide Hospital Research Committee, Early Career Research Fellowship (#17236)

M.T.-A. is the recipient of an Australian Research Council Future Fellowship (FT210100455) and a grant from the Mizutani Foundation for Glycoscience (#250004).

## Author contributions

T.H.C., S.C. and M. T.-A. writing–review and editing;

T.H.C., S.C. and M. T.-A. writing–original draft;

T.H.C., S.C. and M. T.-A. visualisation;

S.C. and R.K. validation;

T.H.C., L.C., A.C. and R.K. methodology;

T.H.C., S.C., L.C., A.C., M.T., H.H., R.K. and M.T.-A. investigation;

T.H.C., S.C., L.C., M.T., H.H., R.K. and M.T.-A. formal analysis;

T.H.C., S.C., L.C. and R.K. data curation;

R. K., E.J.M., D.J.T. and M. T.-A. conceptualisation;

M. T.-A. supervision;

R. K., E.J.M., D.J.T. and M. T.-A. funding acquisition;

E.J.M. and D.J.T. resources;

M. T.-A. project administration.

## Abbreviations

ACN: acetonitrile
AGC: automatic gain control
AGP-1: alpha-1- acid glycoprotein
AUC: area-under-the-curve
CID: collision-induced dissociation
FDR: false discovery rate
glycoPSM: glycopeptide-to-spectrum match
GO: gene ontology
HCD: higher- energy collisional dissociation
HILIC: hydrophilic interaction liquid chromatography
ICU: intensive care unit
Le^x^: Lewis x
LFQ intensity: label free quantitation
ML: machine learning
NCE: normalised collision energy
PGC: porous graphitised carbon
PNGase F: peptide:*N*- glycosidase F
RF: random forest
ROC: receiver operating characteristic
sLe^x^: sialyl Lewis x
SPE: solid-phase extraction
TEAB: triethylammonium bicarbonate
TFA: trifluoroacetic acid
ZIC-HILIC: zwitterionic hydrophilic interaction liquid chromatography.

